# Conserved protein Seb1 that interacts with RNA polymerase II and RNA is a bona fide transcription elongation factor

**DOI:** 10.1101/2024.12.17.628955

**Authors:** Krzysztof Kuś, Soren Nielsen, Nikolay Zenkin, Lidia Vasiljeva

## Abstract

Maturation of protein-coding precursor messenger RNA (pre-mRNA) is closely linked to RNA polymerase II (Pol II) transcription. However, the mechanistic understanding of how RNA processing is coordinated with transcription is incomplete. Conserved proteins interacting with the C-terminal domain of the largest catalytic subunits of Pol II and nascent RNA (CID-RRM factors) were demonstrated to play a role in mRNA 3’-end processing and termination of Pol II transcription. Here, we employ a fully reconstituted system to demonstrate that fission yeast CID-RRM factor Seb1 acts as a *bona fide* elongation factor *in vitro*. Our analyses show that Seb1 exhibits context-dependent regulation of Pol II pausing, capable of either promoting or inhibiting pause site entry. We propose that CID-RRM factors coordinate Pol II transcription and RNA 3’-end processing by modulating the rate of Pol II transcription.

## INTRODUCTION

RNA polymerase II (Pol II) is a multi-subunit machinery that transcribes all protein-coding and many non-coding RNAs. During Pol II transcription, precursor messenger RNA (pre-mRNA) undergoes three essential processing steps: 5’-end capping, splicing, and 3’-end cleavage/polyadenylation to mature into its functional form. Interactions between Pol II and mRNA processing factors are largely mediated by the C-terminal domain (CTD) of the largest subunit of Pol II (Rpb1). The Pol II CTD is composed of the consensus repeats Y_1_S_2_P_3_T_4_S_5_P_6_S_7_ (29 in fission yeast *Schizosaccharomyces pombe* and 52 in humans) where all residues except prolines can be phosphorylated (Buratowski, 2003; Eick & Geyer, 2013; Komarnitsky *et al*, 2000). RNA processing factors show a preference for specific CTD phosphorylation patterns which is a basis for their stage specific recruitment or exchange (Buratowski, 2003; Hsin & Manley, 2012). Among readers of CTD phosphorylation status are proteins with CTD-interacting domain (CID). In addition to CID, many of these proteins contain an RNA-recognition motif domain (RRM) and are named CID-RRM factors (Figure 1A). This protein family including Pcf11, Rtt103 and Seb1 (and its homologues Nrd1 in Saccharomyces cerevisiae; Scaf4/8 in human) regulates key transcriptional events, coordinating the 3’-end processing of nascent transcripts and transition between elongation and termination (Vasiljeva & Buratowski, 2006; Vasiljeva *et al*, 2008; Kim *et al*, 2006; Grzechnik *et al*, 2015; Kamieniarz-Gdula *et al*, 2019; Lemay *et al*, 2016; Wittmann *et al*, 2017; Gregersen *et al*, 2019).

**Figure 1.**
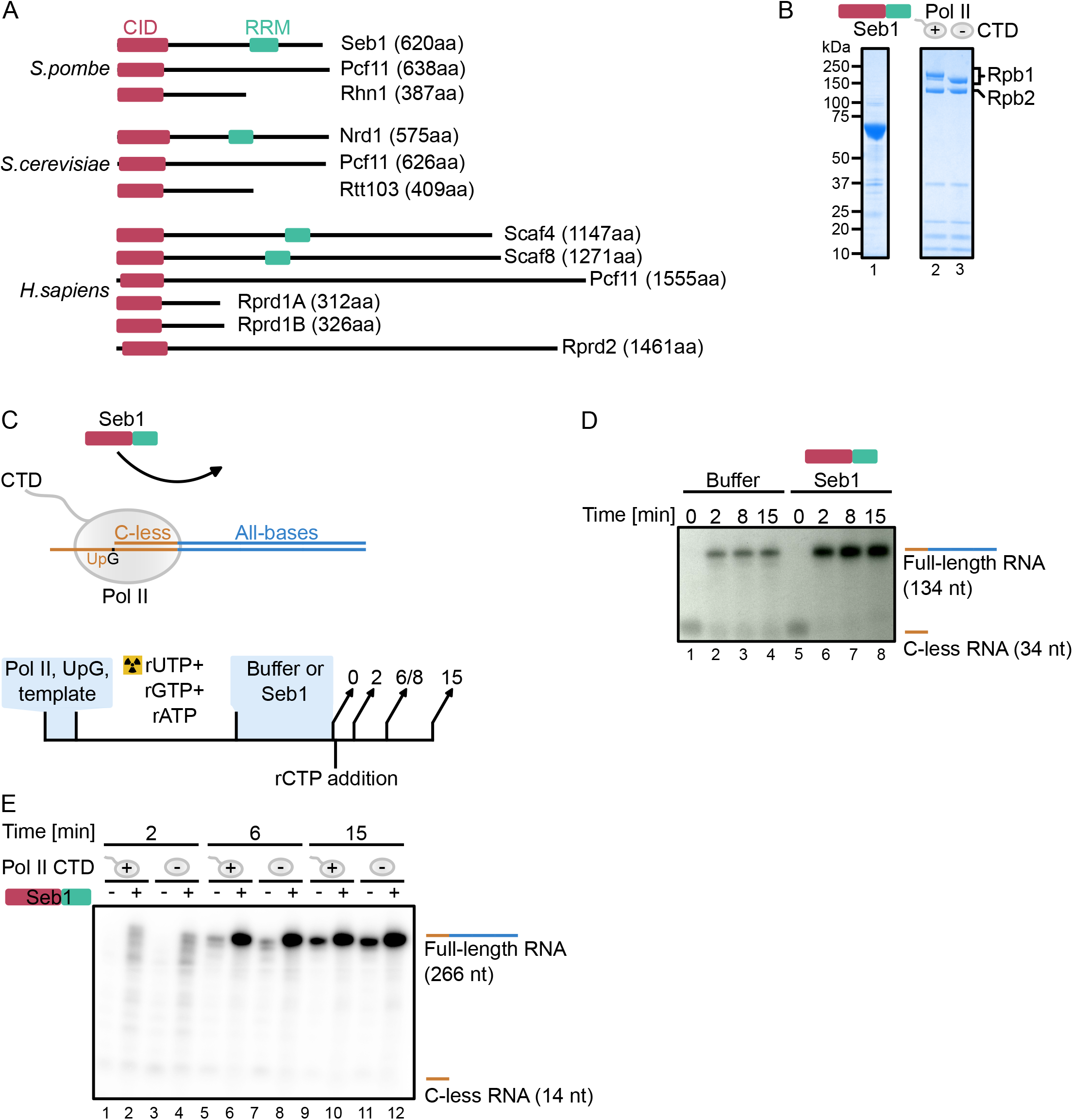
Seb1 stimulates transcriptional output *in vitro*. (A) Seb1 is a conserved protein in eukaryotes. In humans, Seb1 has undergone gene duplication (Scaf4 and Scaf8). This family of proteins contains CTD Interacting Domain (CID) and RNA Recognition Motif (RRM). (B) SDS-PAGE showing exemplary purification of Seb1 or Pol II (+/-CTD). (C) Schematics of *in vitro* transcription which allows multiple rounds of re-initiation. Tailed DNA template with a single-stranded overhang on the 3’ of the template strand has a C-less track allowing stopping elongation complex in the absence of rCTP (upper panel). Transcription is primed by UpG dinucleotide and initiated with α-[^32^P]-UTP, rGTP, rATP. The stopped elongation complex is incubated with or without Seb1 (1.5 µM) and transcription is resumed upon rCTP addition. Products are collected at different time points (lower panel). (D) Seb1 stimulates transcription *in vitro*. Marked C-less RNA is 34 nt whereas extended RNA is 134 nt long. (E) Seb1 stimulatory effects on Pol II do not require CTD. Rpb1 has a TEV-cleavage site which allows efficient removal of CTD (Figure 1B). Marked C-less RNA is 14 nt whereas full-length RNA is 266 nt long.

Transcription termination allows release of RNA and dislodgement of Pol II from the DNA template (Proudfoot, 2016, 2011). This process is tightly linked to the 3’-end processing of nascent RNA – where recognition of polyadenylation signal (PAS) motif by the cleavage and polyadenylation factor (CPF) leads to endonucleolytic cleavage of the transcript (reviewed in (Proudfoot, 2011; Rodríguez-Molina *et al*, 2023; Zhao *et al*, 1999). This cleavage event leaves a downstream, monophosphorylated part of RNA still bound to Pol II and being elongated. It gives a window of opportunity for Xrn2/Rat1/Dhp1 to engage and start degrading nascent RNA which results in Pol II termination (torpedo model) (West *et al*, 2004; Kim *et al*, 2004; Eaton *et al*, 2020; Kuś *et al*, 2023). As alternative, but not mutually exclusive, allosteric model proposes that transcription around the PAS region exerts conformational changes in the Pol II complex aiding termination (Luo & Bentley, 2004; Zhang *et al*, 2015).

The importance of CID-RRM factors is highlighted by their conservation throughout evolution with Seb1 homologues present in humans (Figure 1A). Human Scaf4 and Scaf8 are paralogs that have emerged by gene duplication and share ∼40% amino acid identity between them.

Knockout of Seb1 and both Scaf4/8 is lethal (Wittmann *et al*, 2017; Lemay *et al*, 2016; Gregersen *et al*, 2019). Scaf4 is associated with developmental disorders in humans where one of the alleles is truncated/non-functional. This indicates a haploinsufficiency for Scaf4 and expression of both copies is required for proper transcriptional output (Fliedner *et al*, 2020). Nascent transcriptome analyses upon Scaf8 and Scaf4 depletion indicated that Scaf8 is required for Pol II elongation, whereas Scaf4 is involved in selection of correct PAS by CPF within 3’UTR of the nascent RNA. Seb1 depletion alters transcriptional output, promotes the use of distal polyadenylation sites within 3’ UTRs, and impairs transcription termination (Wittmann *et al*, 2017; Lemay *et al*, 2016). Moreover, Seb1 was linked to the induction of long-lived Pol II pauses during elongation which were proposed to lead to a heterochromatin formation at pericentromeric regions (Parsa *et al*, 2018; Marina *et al*, 2013). These observations were based on the analysis of *seb1-1* mutant that has seven-point mutations (3 changing amino acids and 4 silent mutations that affect translation efficiency without changing the amino acid sequence) (Marina *et al*, 2013).

As mentioned, CID-RRM factors bind to phosphorylated Pol II CTD and Seb1 shows a preference for Ser2P but also can interact with Ser5P whereas Scaf4/8 bind with highest affinity to dual phosphorylated Ser2P, Ser5P (Gregersen *et al*, 2019; Becker *et al*, 2008). This CTD binding bias specifies Seb1 recruitment to the 3’-end of genes where Ser2P CTD phosphorylation peaks (Kecman *et al*, 2018; Gu *et al*, 2013; Lyons *et al*, 2020). Seb1 not only binds to Poll CTD but also is in contact with Pol II core (close to RNA exit channel and Rpb4/7) via region spanning RRM (Kecman *et al*, 2018). These results were obtained without the presence of nucleic acids indicating a direct interaction between Seb1 and Pol II (Kecman *et al*, 2018). In addition, it was previously reported that Seb1 is Rpb7 (part of Pol II stalk) binding protein (Mitsuzawa *et al*, 2003). The putative interface of Seb1-Pol II interaction overlaps with Spt5 (which with Spt4 forms DSIF complex) binding site (Ehara *et al*, 2017; Bernecky *et al*, 2017; Shetty *et al*, 2017; Song & Chen, 2022). In fact, *in vitro* data suggested that Seb1 could outcompete Spt4/5 (Kecman *et al*, 2018). Moreover, Seb1 can interact with RNA close to the promoter region (Lemay *et al*, 2016; Wittmann *et al*, 2017) indicating additional roles beyond termination. Further, both activities of Seb1 (CTD binding and RNA recognition) are required for functional transcription. Mutation compromising CID domain or RRM result in a global transcriptional readthrough in fission yeast (Wittmann *et al*, 2017). Seb1 associates with the CPF complex implying that that CID-RRM factors may directly contribute to mRNA 3’-end processing by CPF leading to changes in PAS usage (Wittmann *et al*, 2017; Lemay *et al*, 2016; Gregersen *et al*, 2019). Notably, Pol II elongation speed has also been linked to PAS selection. Slow Pol II progression favors selection of the more upstream (proximal) PAS, whereas Seb1 depletion favors selection of distant PAS (Pinto *et al*, 2011; Yague-Sanz *et al*, 2020; Geisberg *et al*, 2020; Wittmann *et al*, 2017).

To unambiguously elucidate the role of CID-RRM proteins in Pol II transcription, we employed *in vitro* biochemistry and bioinformatics to investigate how Seb1 affects Pol II elongation. Our results demonstrate that this conserved factor stimulates Pol II transcription in a defined *in vitro* system, while reanalysis of the published data shows that Seb1 might exert pro- or anti-pausing activity on Pol II. Therefore, we conclude that CID-RRM factors are *bona fide* Pol II elongation factors. Seb1 possesses activities that are separated in higher eukaryotes (Scaf4 and Scaf8) and our results provide better understanding of how coupling between transcription and RNA processing is controlled by the CID-RRM factors.

## RESULTS

### Seb1 is an elongation factor *in vitro*

As Seb1 and its homologues are key to Pol II transcription, we employed a reductionist *in vitro* system to uncover its direct contribution to transcription. To this end, we expressed Seb1 in *E. coli* as N-terminal His-tag fusion. Seb1 was purified using affinity chromatography, followed by gel filtration (Figure 1B, lane 1). Pol II was obtained from a native source (*S. pombe*) using a strain with a 3xFLAG-tag on Rpb9. Polymerase was purified using affinity (Flag-M2 agarose resin), followed by ion exchange chromatography (Q column) (Figure 1B, lane 2). To test how Seb1 contributes to transcription, we employed two complementary experimental *in vitro* strategies. In the first approach, we used a “tailed” DNA template with a single-stranded overhang on the 3’ of the template strand, thus allowing factor-less transcription initiation with UpG primer. The transcribed sequence started with C-less cassette allowing formation of a halted elongation complex in the absence of rCTP and, thus, synchronisation of transcription (Figure 1C, 1D, lanes 1 and 5, and Table S1). The halted elongation complex can then be incubated with factors of interest (Seb1), and transcription resumed by rCTP addition. This setup allows multiple rounds of transcription (Figure 1C). Addition of Seb1 to the stalled elongation complex had strong stimulatory effect on further transcription (Figure 1D, compare lanes 2 and 6). This is surprising taking into account previously published observation suggesting that Seb1 is needed to promote Pol II pausing (Parsa *et al*, 2018).

Given that Seb1 interacts with both the CTD and core enzyme of Pol II, we investigated whether the CTD is essential for this stimulatory effect. To achieve this, we used a strain with a TEV protease cleavage site upstream of the CTD allowing proteolytic removal of this domain. Pol II-TEV-CTD was purified, treated with either TEV protease or buffer, and further subjected to gel filtration (Figure 1B, lane 3). Next, the experiment was performed using Pol II with or without this repetitive region (Figure 1B and E). Both polymerases (+/-CTD) showed comparable activity (Figure 1E, compare lanes 5 and 7). Again, we noted that the presence of Seb1 led to increased amounts of full-length RNA (Figure 1E, compare lanes 5 and 6). Although Seb1 has a CID domain, we have not observed differences in the stimulation of Pol II with or without CTD (Figure 1E, compare lanes 6 and 8). Therefore, we conclude that Seb1 does not require CTD to enhance transcription in this setup.

As we could not exclude the possibility that Seb1 dislodge Pol II from the template (terminate Pol II) and facilitate another round of transcription, we utilised an alternative *in vitro* system, using assembled elongation complexes, immobilized on streptavidin beads via biotin on the non-template DNA strand (Figure 2A). Assembled elongation complexes are indistinguishable from native elongation complexes and exclude possibility of reinitiation of transcription (Figure 2B). Consistently, Seb1 stimulated RNA extension in concentration-dependent manner and resulted in fewer pauses (Figure 2B, compare lanes 6 and 10). Pausing can lead to backtracking, when Pol II shifts backwards with RNA 3’-termini leaving the catalytic center via secondary channel. To explore whether some of the pauses that are controlled by Seb1 represent backtracked Pol II, we tested their sensitivity to TFIIS, which stimulates an intrinsic Pol II activity to cleave backtracked RNA thus reinstating the 3’-end in the active center and reactivating elongation complex for further RNA extension (Noe Gonzalez *et al*, 2021; Sheridan *et al*, 2019; Izban & Luse, 1992; Reines, 1992). We expressed the truncated form of TFIIS in *E. coli* and purified it to homogeneity (Figure 2C, left panel). We then evaluated the impact of increasing amounts of TFIIS on the pausing pattern and we observed accumulation of the full-length RNA accompanied by the simultaneous disappearance of the most prominent pauses, indicating that the paused elongation complexes are mainly backtracked ones. This observation led us to investigate whether Seb1 might resolve TFIIS-sensitive pauses, albeit with slower kinetics. To test this possibility, we performed transcription reactions using higher nucleotide concentrations and collected samples after 5 or 10 min. Only in the presence of Seb1, pausing was decreased leading to more full-length product (Figure 2D, compare lanes 3, 4 with lanes 5, 6). These results suggest that Seb1 may inhibit Pol II backtracking, possibly by interacting with the transcript behind the elongation complex and blocking its reverse movement. Collectively, Seb1 exhibits properties of an elongation factor that can control pausing *in vitro*.

**Figure 2.**
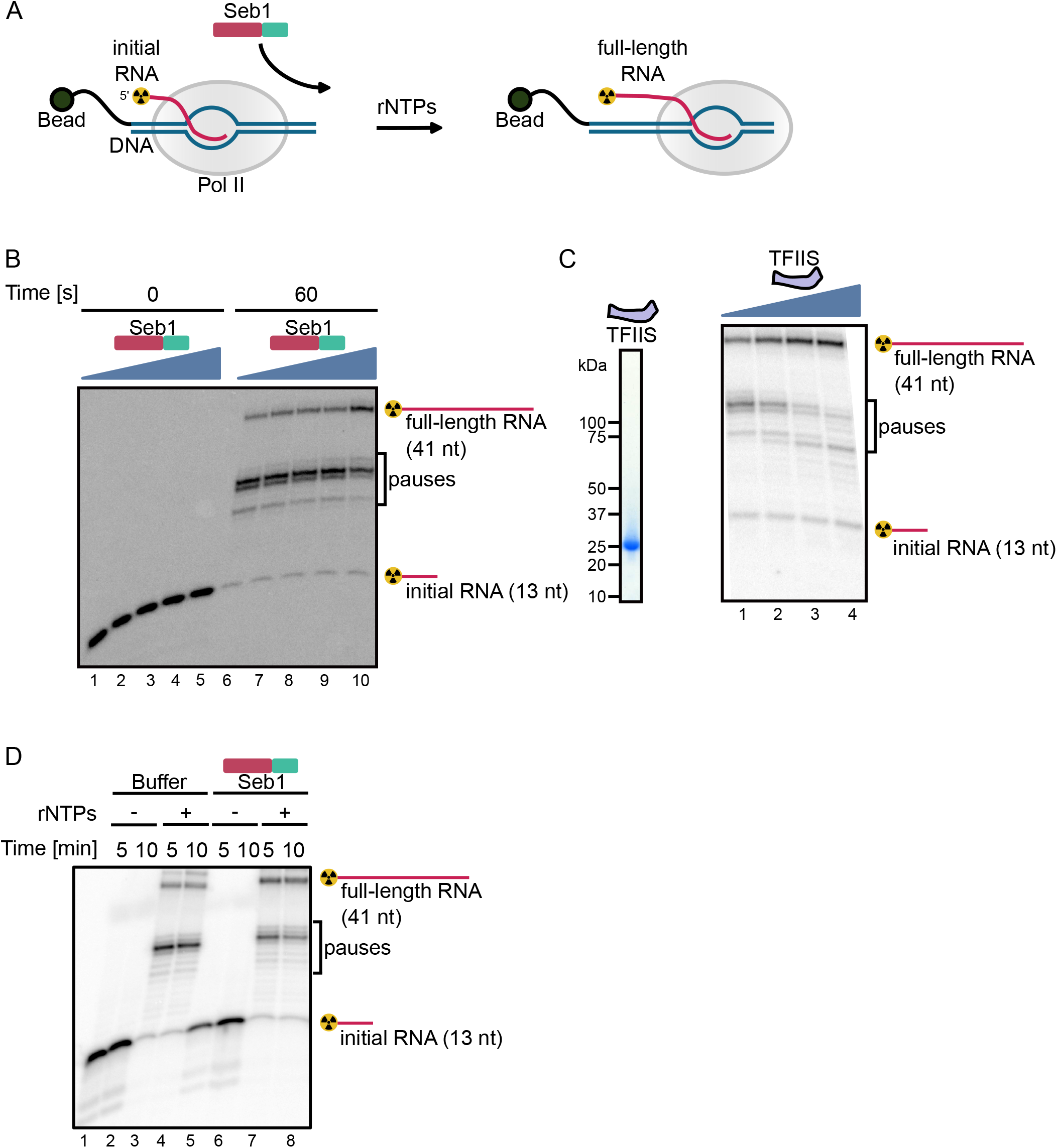
Seb1 affects pausing *in vitro*. (A) Schematics of *in vitro* assay with Pol II transcriptional elongation complex immobilised on streptavidin beads via biotin on the DNA template strand. RNA is ^32^P labelled on 5’-end. (B) Seb1 decreases pausing in concentration-dependent manner. Initial RNA has a length of 13 nt and full-length one 41 nt, respectively. Transcription complexes were incubated with 0, 0.5, 1, 2 or 4 µM Seb1 and transcription was initiated with 100 µM rNTPs and 10 mM Mg^2+^. (C) The left panel shows SDS-gel of purified TFIIS (*S. pombe* TFIIS_115-293_). Pausing pattern changes when TFIIS_115-293_ (0, 25 nM, 75 nM or 240 nM) is present during 30 s transcription (right panel) indicating backtracking of the elongation complex in these positions. (D) Seb1 aids in overcoming TFIIS-sensitive pauses. Transcription complexes were incubated with or without 800 nM Seb1 and transcription was initiated with 0.9 mM rNTPs and 5 mM Mg^2+^.

### Seb1 can lead to different Pol II pausing outcomes *in vivo*

Seb1 has been associated with controlling Pol II pausing *in vivo* (Parsa *et al*, 2018). It was demonstrated that *seb1-1* mutant exhibits defects in heterochromatic silencing at pericentromeric region (Marina *et al*, 2013). These effects were linked to compromised recruitment of NuRD-related chromatin-modifying complex SHREC and lack of long-lived Pol II pauses in *seb1-1* mutant as evaluated by native elongating transcript sequencing (NET-seq). It was postulated that Seb1 controls Pol II pausing which serves as a signal for heterochromatin nucleation (Marina *et al*, 2013; Parsa *et al*, 2018). In line with these findings, we decided to revisit NET-seq data to explore different classes of pausing in the *seb1-1* allele (Parsa *et al*, 2018). Interestingly, the additional analyses of NET-seq data had revealed that mutations in Seb1 can lead to either positive or negative effects on Pol II pausing in the promoter region (Figure 3A). This analysis was performed genome-wide, focusing on protein-coding genes not containing other genes 250 bp upstream TSS or 250 bp downstream PAS. We calculated the promoter pausing index for the genes with NET-seq signal (refer to Methods). Using this approach, we found ∼250 genes that had increased pausing in the promoter region in the *seb1-1* strain and we plotted it as a metaprofile (Figure 3B, left panel). As a comparison, we present a group of genes that showed a drop in the pausing index in the promoter region (Figure 3B, right panel). Nevertheless, in both gene classes, there is an additional peak around the PAS region, suggesting that Pol II accumulates at 3’-end for both gene groups in *seb1-1* strains.

**Figure 3.**
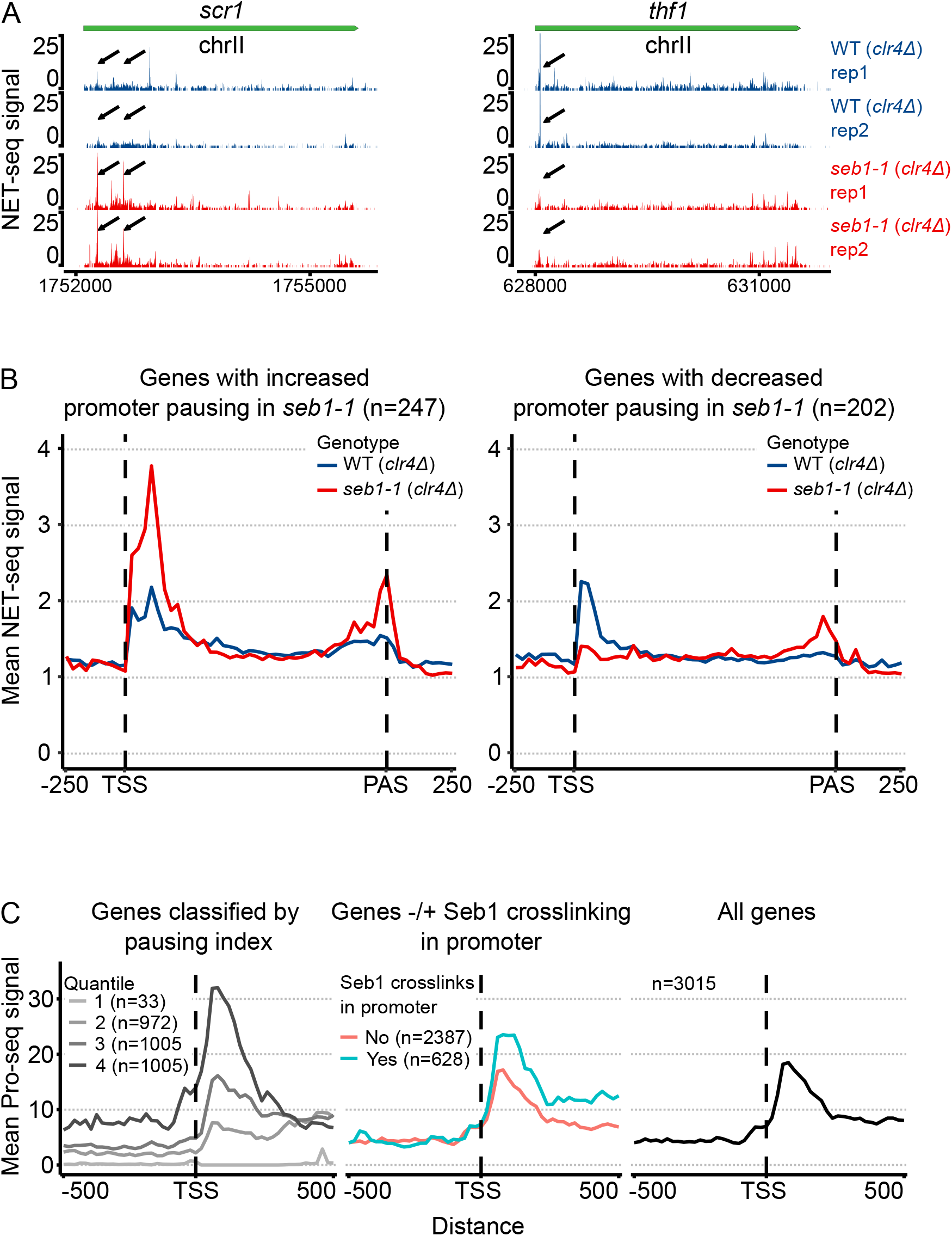
Seb1 correlates with pausing *in vivo*. (A) Seb1 might act as a positive or negative regulator of the Pol II pausing. Two example genes showing the opposite effect on pausing in *seb1-1* cells. (B) Metaprofile for the subset of genes exhibiting increased or decreased pausing index in the promoter region (NET-Seq) for reference and *seb1-1* strain. (C) Association between Pol II pausing and genes with Seb1 crosslinking in the promoter region (middle panel). The left panel depicts genes split into quantiles according to the pausing index and the right one is the average of all genes considered.

To further explore the correlation between Seb1 and pausing we decided to stratify genes into two categories depending on whether the gene had Seb1 crosslinking in the promoter as found in our Seb1 PAR-CLIP data (Wittmann *et al*, 2017). Next, we plotted the mean Pol II PRO-seq signal and noted that on average genes with Seb1 associated with RNA at the 5’-end of transcription show elevated pausing close to TSS (Wittmann *et al*, 2017; Booth *et al*, 2016). As a reference, we present metaprofiles for all genes or genes split into 4 quantiles depending on the pausing index (Figure 3C, left and right panels). We propose that Seb1 can promote pausing at TSS or alternatively it might facilitate pause release of Pol II at genes that have high level of paused Pol II. Taken together, we suggest that Seb1 is an elongation factor that can contribute to control of Pol II pausing in context-dependent manner.

## DISCUSSION

The 3’-end processing and termination of Pol II generates functional RNA molecules and allows polymerase recycling. CID/CID-RRM factors are central to the coordination of the last transcriptional steps. Here, we focused on Seb1 as a model for the CID-RRM factors and tested its functions in the defined *in vitro* systems and observed that Seb1 can promote Pol II transcription. Moreover, we provide evidence that Seb1 prevents TFIIS-sensitive pauses *in vitro*. We also uncovered that Pol II CTD is not necessary for Seb1 stimulation which points towards this repetitive region functioning as a recruitment platform for CID-RRM factors that can be less important in the *in vitro* settings where Seb1 amounts are saturating. Seb1 RRM is in contact with the Pol II core close to the RNA channel and RNA binding of the nascent RNA might prevent of long-lived backtracked pauses (Kecman *et al*, 2018). This may happen via binding the transcript emerging from Pol II RNA-exit channel and physical blocking of backward movement of the elongation complex, i.e. backtracking. An allosteric effect on elongation complex can also take place, in a manner of antitermination factor p7 action on bacterial RNA polymerase, that biases elongation complex away from the backward movement (Yuzenkova *et al*, 2008).While Seb1 potentially can outcompete Spt5 for Pol II binding, we can envision a possibility of the co-existence of these factors in the elongation/initiation phase (Kecman *et al*, 2018). In fact, modular structure of Spt5 would be compatible with re-arrangements of interactions within Pol II core without Spt5 full dissociation from Pol II (Kuś *et al*, 2023; Zeng *et al*, 2024; Yanagisawa *et al*, 2024).

To correlate our *in vitro* findings, we decided to revisit publicly available data to provide correlations between Seb1 binding and pausing *in vivo* (Wittmann *et al*, 2017; Parsa *et al*, 2018; Booth *et al*, 2016). Although Seb1 was proposed to induce long-lived pauses *in vivo*, we observed that for a subset of genes this protein has the opposite effects (Parsa *et al*, 2018). Therefore, Seb1 might exhibit context-dependent control of Pol II pausing. In addition, the accumulation of pauses close to the PAS in *seb1-1* mutant is compatible with the idea that Seb1 might modulate CPF cleavage and when this process is compromised Pol II accumulates around this region. Furthermore, we noted that genes with Seb1 crosslinking in their promoter regions show higher pausing propensity compared to genes lacking Seb1 PAR-CLIP signals. This correlation suggests two possible interpretations: either Seb1 directly promotes pausing potentially via binding to RNA, or alternatively, Seb1 preferentially targets already-paused genes to regulate their transition into elongation.

It might appear counter-intuitive that Seb1 or Scaf4/8 might act as termination and elongation factors. Nevertheless, the dualism of action for transcriptional factors has been observed before including Spt5 which can promote pausing at TSS or act as a positive elongation factor (Wada *et al*, 1998; Hu *et al*, 2021; Bourgeois *et al*, 2002). In bacteria, NusG and NusA display opposing effects on termination depending on the external cues (Mooney *et al*, 2009; Wells *et al*, 2016). This functional plasticity may extend to Seb1, where its functionality might be modulated by the binding partners (RNA or CTD) and post-translational modifications particularly phosphorylation of its multiple residues could serve as a molecular switch between pro- and anti-termination states (Swaffer *et al*, 2018). Alternatively, the composition of the complexes associated with Pol II might dictate which role Seb1 takes in each transcriptional step. Therefore, integration of multiple signals might be required for Pol II to undergo 3’-end processing and termination. Seb1 could be a nexus for such regulation as it can form multivalent interaction with Pol II (CTD and core), nascent RNA but also with CPF complex. Although, we provide *in vitro* data and *in vivo* correlates that Seb1 can act as a positive elongation factor, further research is required to evaluate the impact of Seb1 on the transcriptional elongation. This could be achieved with rapid Seb1 depletion and measurements of nascent transcription.

In summary, we propose that Seb1 can control Pol II pausing and act as an elongation factor beyond involvement in 3’-end processing/termination. These functionalities appear to be separated in higher eukaryotes, where gene duplication created paralogs (Scaf4 and Scaf8) that possess overlapping but also unique functions. Differences in genome organisation and length of the genes could have been a driving force for a separation of Seb1 homologues functions. Scaf4 seems to resemble more Seb1 3’-end functionalities and both Scaf4/8 act as positive elongation factors (anti-terminators). Overall, Seb1 might function as a context-dependent regulator of Pol II progression ascertaining that 3’-end processing and transcription termination would generate functional RNA molecules.

## METHODS

### Protein purification

Pol II was purified in a similar manner as previously described (Kecman *et al*, 2018). Briefly, *S. pombe* yeast strain with endogenously tagged Rpb9-3xFLAG (and TEV cleavage site before CTD of Rpb1) was grown in YES media (genotype: h+, leu1-32, ura4-Δ18, ade6-M216; his3Δ::1; Rpb9-3xFLAG::kanMX; Rpb1-TEV-CTD - a TEV cleavage site was inserted upstream the CTD), harvested and stored at −80 °C until day of purification. Cells were lysed in a freezer mill (SPEX SamplePrep) in liquid nitrogen and mixed with lysis buffer (50 mM Tris-HCl pH 7.5, 150 mM NaCl, 10% glycerol, 0.5% Triton X-100, 0.5 mM DTT, 0.5 mM MgCl_2_, 0.5 mM Mg(OAc)_2_) supplemented with proteinase inhibitor cocktail (cOmplete, EDTA-free Protease Inhibitor Cocktail, Roche). Lysate was cleared using centrifugation at 40000 x g for 20 min and incubated with α-FLAG beads (M2 agarose gel, Sigma) for 1.5 h in cold room. Beads were washed three times with W1 buffer (50 mM Tris-HCl pH 7.5, 1 M NaCl, 1 M urea, 10% glycerol, 0.5% Triton X-100, 0.5 mM DTT, 0.5 mM MgCl_2_, 0.5 mM Mg(OAc)_2_) and three times with W2 (as W1 but 150 mM NaCl, no urea, 0.05% Triton X-100). Next, Pol II was eluted with 5 mL of 2.5 mg/mL 3xFLAG-peptide (Sigma), and mixed with 20 mL of Q_A buffer (50 mM TRIS pH 7.7, 5 mM NaCl, 10% glycerol, 0.5 mM MgCl_2_, 0.5 Mg(OAc)_2_, 1 mM β-mercaptoethanol). Protein eluate was then applied to an ion exchange chromatography column (2 x 1 mL or 1x 5 mL HiTrap Q HP, GE Healthcare) equilibrated with Q_A buffer. The column was washed with several column volumes of 8% buffer Q_B (as Q_A but 2 M NaCl). Protein was eluted with a gradient of Q_B buffer (up to 40%). The buffer was exchanged to storage buffer (20 mM HEPES pH 7.5, 150 mM NaCl, 0.5 mM MgCl_2_, 0.5 mM Mg(OAc)_2_, 1 mM β-mercaptoethanol) and proteins snap-frozen and kept at −80 °C till use. To remove CTD, Pol II with TEV cleavage site was incubated with AcTEV (Invitrogen) at room temperature. Pol II with or without CTD was subjected to gel filtration on Superose 6 3.2/300 (GE Healthcare), aliquoted and stored −80 °C for further experiments.

Seb1 (full-length, with C-terminal His-tag, in pET41a plasmid) was expressed in Rossetta *E. coli* strain at 37 °C using IPTG induction for 4 h. Cells were harvested by centrifugation at 4 °C and frozen. Cell pellets were resuspended in NiTA buffer (50 mM Tris-HCl pH 7.7, 600 mM NaCl, 5 mM imidazole, 1 mM β-mercaptoethanol) with proteinase inhibitor mix. Cells were lysed using French Press and phenylmethylsulfonyl fluoride (PMSF) was added 1 mM final concentration. Lysates were spun at 20000 x g, 4 °C for 30 min, filtered and incubated with Ni-NTA Agarose (Qiagen, 1 mL slurry per 2 L of cell culture) for 1h. Beads were washed with several column volumes of NiTA buffer and protein was eluted with 500 mM imidazole. Further protein was centrifuged at 10000 x g for 10 min and subjected to gel filtration on Superdex 200 10/300 (GE Healthcare) equilibrated with GF buffer (20 mM HEPES pH 7.6, 150 mM NaCl, 1 mM DTT, 0.5 mM MgCl_2_ and 0.5 mM Mg(OAc)_2_). Protein was snap-frozen and kept at −80 °C till use.

Truncation of TFIIS (residues 115-293 aa of *S. pombe tfs1* gene, codon optimised for *E. coli*, as N-terminal His-tag, with TEV cleavage site) was purified in a similar manner as Seb1 except nickel affinity was done on HisTrap HP (GE Healthcare), followed by ion exchange (HiTrap SP HP, GE Healthcare) and with final gel filtration on HiLoad 16/60 Superdex, 75 PG. Proteins were snap-frozen and kept at −80 °C.

### *In vitro* transcription

Tailed DNA templates were prepared using PCR, cleaved with BsaI enzyme (New England Biolabs), dephosphorylated with antarctic phosphatase (New England Biolabs), ligated with a single-stranded phosphorylated oligo (to create a 3’-end overhang on template strand) and purified using PCR purification kit (Qiagen). Sequences are listed in Table S1. Typically, starting 20 µl reaction containing tailed DNA template (125-170 ng), 1 µl UpG (7.5 mM dinucleotide primer, Jena Bioscience), Pol II (250-400 ng) and 5 µl rNTPs (50 µM rATP, 50 µM rGTP, 5 µM rUTP with added 1 µl hot α-^32^P-rUTP 0.37 MBq/µL, Hartmann Analytic without rCTP), RNAsin (0.5 µl, Promega) was incubated in TB1 buffer (TB1: 37.5 mM HEPES pH 7.5, 150 mM KCl, 0.375 mM DTT, 0.75 mM MnCl_2_, 0.75 mM MgCl_2_, 0.75 mM Mg(OAc)_2_, 9% glycerol). This allowed transcription of C-less segment and reactions were carried for 25-30 min at room temperature. Proteins (1/5 or 1/6 of reaction volume, final concentration indicated in the figure) were incubated with the complex (10-15 min) and rCTP (10-12.5 µM) was used to resume transcription. Samples were collected at indicated time-points and stopped with addition of 0.2 mg/mL proteinase K, 12 mM EDTA, 0.2% SDS. RNA products were separated on 6% or 8% PAGE-UREA gel and visualised on FLA-7000 phosphoimager (Fujifilm) or autoradiography film (Amersham Hyperfilm MP).

Artificial elongation complexes were assembled as described by us earlier (Nielsen & Zenkin, 2013; Heo *et al*, 2021). Briefly, reactions were assembled in a stepwise manner in TB buffer (20 mM Tris, 40 mM KCl, 200 µM EDTA, pH adjusted to 7.9 with HCl). First, for final standard 10 µl reaction, ∼2.5 pmol of 5’ radioactively labelled RNA was annealed with 1 pmol of template DNA at 45 °C and slowly cooled down to room temperature. Next, 0.3-0.4 pmol of Pol II was added to the mix and kept for 10 minutes at 30 ºC, followed by same incubation with 10 pmol of non-template DNA (Table S1, T-DNA, NT-DNA, RNA13 labelled with ^32^P on 5’). Complex was immobilised on equilibrated streptavidin beads (Streptavidin Sepharose HP, Cytiva, 6 µl per reaction) as previously reported (Nielsen *et al*, 2013). Beads were washed once with TB, once with W500 (20 mM TRIS, 500 mM NaCl pH adjusted to 7.9 with HCl) and again 3 times with TB. After that the complex was incubated with buffer or protein at concentration specified in the figure legend for at least 5 min at 30 ºC. Transcription was started with the addition of rNTPs/Mg^2+^ at concentration indicated in figure legend and samples were collected at times specified (reaction stopped with equal volume of formamide containing: 20 mM EDTA, 7 M urea, 100 μg/mL heparin, 0.02% bromophenol blue, 0.03% xylencyanol in formamide). Products were resolved on 10% UREA-PAGE gel and visualised on FLA-7000 phosphoimager (Fujifilm). Reactions were performed at least twice (with same or different templates).

### Bioinformatics

For the analysis only coding genes were considered and it was required that gene does not have another transcription unit annotated on the same strand in 250 bp before TSS or after PAS (n=3190). Publicly available NET-Seq data (accession: GSE114540) in the bigwig format was used to calculate pausing index. Only genes longer than 900 bp and having signal >10 in gene body were used. The pausing index was defined as ratio of the signal in 300 bp downstream TSS, divided by read density in in gene body (region TSS+300 bp to PAS-300 bp). To select genes with pausing index increased in *seb1-1* strain, we required that the trend is consistent between the replicates with a difference of pausing index at least 0.5. Most likely our analysis underestimates the number of genes with increased/decreased pausing as we limited our analysis to the subset of genes. Metaprofiles were prepared for each replicate with deepTools (Ramírez *et al*, 2016) averaged and plotted in R package (Ihaka & Gentleman, 1996).

To correlate Seb1 RNA binding and pausing, we used Clip data (accession: GSE93344) (Wittmann *et al*, 2017) and PRO-Seq data (accession: GSM1974985) (Booth *et al*, 2016). The pausing index was defined as above except genes with any signal in gene body and longer than 600 bp were considered (n=3015). Gene body was defined as a region between TSS+300 bp till PAS. Next, genes were assigned according to presence of the Seb1 cross-linking in the promoter. Metaprofiles (all genes average, quantiles and genes with/without Seb1 crosslinks in the promoter region) were plotted in R environment.

## Acknowledgements

This work was supported by a Wellcome Trust Senior Research fellowship and BBSRC grants to LV (WT106994/Z/15/Z and BB/Y00194X/1) and Wellcome Trust Investigator Award to N.Z. (217189/Z/19/Z).

## Author Contributions

K.K and L.V. conceived and designed the experiments and wrote the manuscript. K.K. performed experiments and bioinformatics correlations. S.N. contributed to the establishment of *in vitro* transcription assay in N.Z. laboratory. N.Z. provided *in vitro* transcription expertise. All authors edited the manuscript.

